# cGAS/STING and NLRP3 cooperatively activate CD8+ T cell-mediated anti-tumor immunity in colorectal cancer

**DOI:** 10.1101/2023.08.22.554371

**Authors:** Courtney Mowat, Daniel Schiller, Kristi Baker

## Abstract

Colorectal cancer (CRC) is a highly prevalent and deadly disease that is largely refractory to immunotherapy. The only CRC subset that responds to these therapies is characterized by prevalent microsatellite instability (MSI), extensive CD8+ T cell infiltration and high expression of innate immune signaling pathways. Endogenous activation of the cGAS/STING pathway is essential for the CD8+ T cell antitumor response in MSI CRCs, suggesting that activating it in other CRCs could boost immunotherapy response rates. We show that cGAS/STING signaling can be enhanced by costimulation of the NLRP3 inflammasome and that dual stimulation increases CD8+ T cell-mediated antitumor immunity in both MSI and non-MSI CRCs. The ability of NLRP3 to boost cGAS/STING signaling was specific and did not occur with activation of other innate immune pathways such as AIM2 or TLRs. Cooperativity between cGAS/STING and NLRP3 proceeded via a positive feedback loop that was inflammasome-independent and required early crosstalk between the signaling mediators and regulation of their gene expression. Notably, increased cGAS/STING signaling enhanced CD8+ T cell activation when in conjunction with anti-PD1 immunotherapy, suggesting that signaling via NLRP3 could further boost this response and render otherwise resistant CRC susceptible to immunotherapy.

**Significance:** Innate immune signaling pathways cooperatively regulate CD8+ T cell-mediated antitumor immunity in both hot and cold tumors. In addition to serving as predictive biomarkers, these pathways can be therapeutically targeted to increase response rates to immunotherapy while minimizing undesirable adverse events.

## Introduction

Continuous exposure of the intestine to foreign lumenal antigens forces the mucosal immune system to be tolerogenic to avoid constitutively inflammation. Unfortunately, such immune tolerance generates a fertile ground for the development of colorectal cancers (CRCs). The majority of CRCs are induced by *APC* mutations and develop extensive chromosomal instability (CIN) due to rearrangements and missegregation.^1,2^ These cancers have a low tumor mutation burden (TMB), high rates of invasiveness and metastasis and possess very few infiltrating immune cells.^3^ In contrast, the 15% of CRCs induced by hypermethylation of the *MLH1* promoter develop widespread point mutations and microsatellite instability (MSI) due to loss of DNA mismatch repair.^3–5^ These CRCs have a much more favorable prognosis despite their high TMB and, notably, contain abundant CD8+ tumor infiltrating lymphocytes (TILs). While this high immunogenicity of MSI CRCs was initially attributed solely to the high levels of neoantigens resulting from the hypermutable genomes, it is now recognized that this is insufficient to drive successful antitumor immunity. Indeed, cancers in other tissue locations such as the endometrium, pancreas and breast that have MSI do not experience a more favorable prognosis despite their high neoantigen load.^6–8^

An additional component necessary for successful antitumor immunity is activation of specific costimulatory pathways that facilitate recruitment and activation of CD8+ T cells. One such pathway recognized to be critical for antitumor immunity is the type I interferon (IFN) pathway that produces IFNA/B.^9^ These cytokines are essential for maximal CD8+ T cell development, differentiation, localization and activation. While production of cytokines such as IFNA/B in the tumor microenvironment (TME) is often attributed to immune cells, epithelial cells are also powerful modulators of the immune system. This is arguably most strongly manifested in the intestine where intestinal epithelial cells are key regulators of local mucosal immunity and serve as a bridge between the microbial and dietary components of the lumen and the diverse population of mucosal immune cells residing in the intestine.^10^ Key molecules involved in this process are the pattern recognition receptors (PRRs) that recognize conserved microbial and innate structures known, respectively, as pathogen-associated molecular patterns (PAMPs) and damage-associated molecular patterns (DAMPs).^11,12^

The contribution of one PRR pathway to antitumor immunity has increasingly been investigated especially in the context of MSI CRCs. The cGAS/STING cytosolic dsDNA-sensing pathway is constitutively activated in MSI CRCs due to endogenous DNA damage. This activates type I IFN signaling within the cells that directly promotes recruitment and activation of CD8+ TILs.^13^ However, rarely is a single PRR activated in isolation since most PAMPs and DAMPs occur within cells or organisms containing many other potential PRR ligands. Given that the outcome of any PRR activation will depend upon crosstalk with other activated pathways, it is important to determine how activation of other PRRs will affect the outcome of cGAS/STING signaling in ways that promote or dampen the antitumor immune response. One pathway of note that is likely to have a considerable impact on antitumor immunity is the NLRP3 inflammasome pathway which responds to various stress-associated PAMPs and DAMPs to trigger inflammation. In the context of cancer, this can either promote tumor progression and inhibit effective priming of TILs, thereby decreasing overall antitumor immunity, or it could release cytokines that activate CD8+ T cells, enhancing effective tumor clearance.^14,15^ How NLRP3 contributes to cGAS/STING signaling in CRC cells themselves remains unknown but could provide important information for understanding a CRC patients’ prognosis and likely responsiveness to immunotherapies.

## Results

### NLRP3 and cGAS/STING expression and activation cooccur in human MSI CRC and correlate with CD8+ TILs

WE first approached this question by mining the PanCancer Atlas CRC dataset in The Cancer Genome Atlas (TCGA) database to determine the overall expression levels of the cGAS/STING and NLRP3 mediators. We noted that MSI CRCs have significantly higher mRNA expression of both *STING* and *NLRP3* (**Fig. 1A**), as well as of many of the canonical products associated with each signaling pathway (**Fig. 1B** and Supplementary Fig. 1A). Furthermore, we identified a significant positive correlation between the transcript expression of *STING* and *NLRP3*, as well as between either *STING* or *NLRP3* and the cytotoxic T cell marker *CD8A* in CRCs (**Fig. 1C**). While the correlation of *STING* and *CD8A* was not unexpected, the correlation between proinflammatory *NLRP3* and *CD8A* was somewhat surprising. However, since the RNAseq data in TCGA is derived from whole tumor tissue, we did not know if the expression of *STING* or *NLRP3* was in the tumor cells themselves or in the immune cells infiltrating the tumor. This is important because the higher numbers of immune cells infiltrating MSI compared to CIN CRCs could provide the false impression that MSI CRCs themselves are directly orchestrating this relationship when it is being indirectly controlled by the TME.

**Figure 1.**
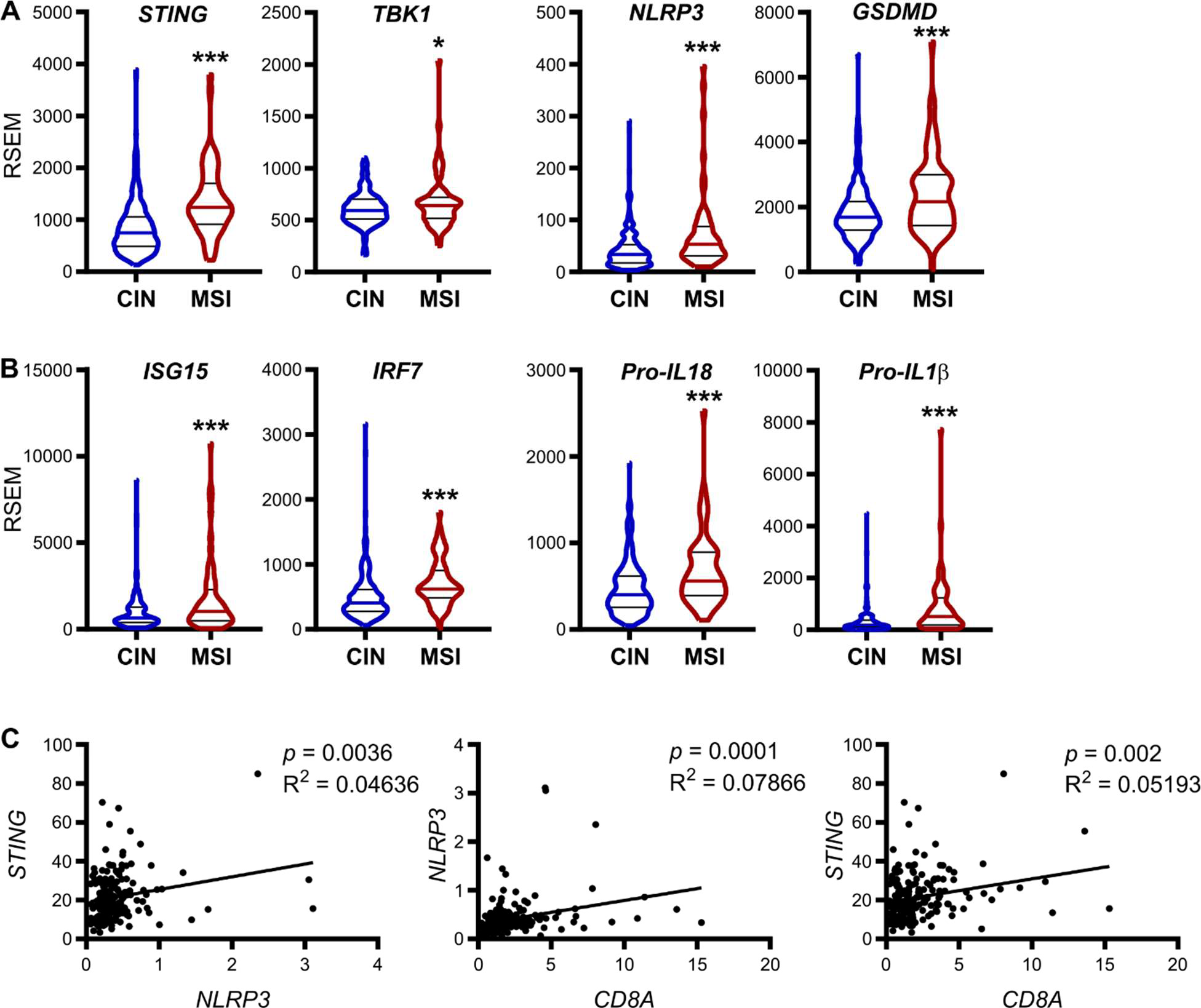
Human MSI CRCs have significantly higher expression of *NLRP3* and *STING* compared to CIN mutated CRCs and the two pathways are highly correlated. **(A)** RNA sequencing for the expression of *cGAS/STING* and *NLRP3* signaling pathway mediators in human CIN and MSI colorectal cancers from the PanCancer Atlas CRC dataset on the TCGA database. **(B)** Expression of major products of the *cGAS/STING* and *NLRP3* inflammasome signaling pathways. Y-axis RNAseq units are in RNA-Seq by Expectation-Maximization (RSEM) Linear regression (C.I. = 95%) was performed with the Pearson correlation R^2^ and *p*-values reported on each graph. **(C)** Correlation of expression of between *STING*, *NLRP3* and *CD8A* expression in all CRCs of the dataset. Panels A-B: \**p* = 0.05, \*\*\**p* < 0.001.

To better tease apart the contributions of the tumor and immune cells, we generated MSI and CIN variants of the mouse MC38 CRC cell line by deleting *Mlh1* and mutating the CIN-associated gene *Kras*, respectively.^16^ Each variant was then subcutaneously injected into the flank of immunocompetent C57BL/6 mice. We confirmed immunohistochemically and by flow cytometry that the *Mlh1^-/-^* MSI tumors contained greater numbers of CD8+ TILs and expressed higher levels of STING than the CIN tumors and thus phenocopied the main characteristics of MSI CRC in patients (**Fig. 2A** and **B**). When we examined NLRP3 and phospho-STING (pSTING) expression in the tumor cells, we noted that each was negatively correlated with tumor mass regardless of whether the tumors were MSI or CIN (**Fig. 2C**). This suggested a role for both of these signaling pathways in anti-tumor immunity. Interestingly, we found a very strong correlation between the expression of pSTING and NLRP3 within the tumor cells themselves, suggesting that activation of these pathways might be linked (**Fig. 2D**). This hypothesis was strengthened when we separately isolated the epithelial and immune cells and assessed RNA expression of known downstream products for each pathway. We found that the NLRP3 inflammasome product pro-*IL1B* was correlated not only to *Nlrp3* expression but also to *Sting* expression in the CRC cells (**Fig. 2E**). Similarly, expression of the STING pathway product *Ccl5* was correlated not only to CRC *Sting* expression but also to *Nlrp3* expression. Given that we have shown CCL5 to be critical for recruitment of CD8+ T cells into the tumor microenvironment (TME), we examined the relationship of *Sting* and *Nlrp3* expression to *Cd8a* expression in the tumors.^16^ This showed that both *Sting* and *Nlrp3* expression in the CRC cells are correlated with *Cd8a* within the TILs but that *Sting* and *Nlrp3* expression in the TILs themselves are not (Supplementary Fig. 1B and C). These data indicate that these two PRRs each contribute to promoting CD8+ T cell-mediated antitumor immunity and that they may work together to accomplish this.

**Figure 2.**
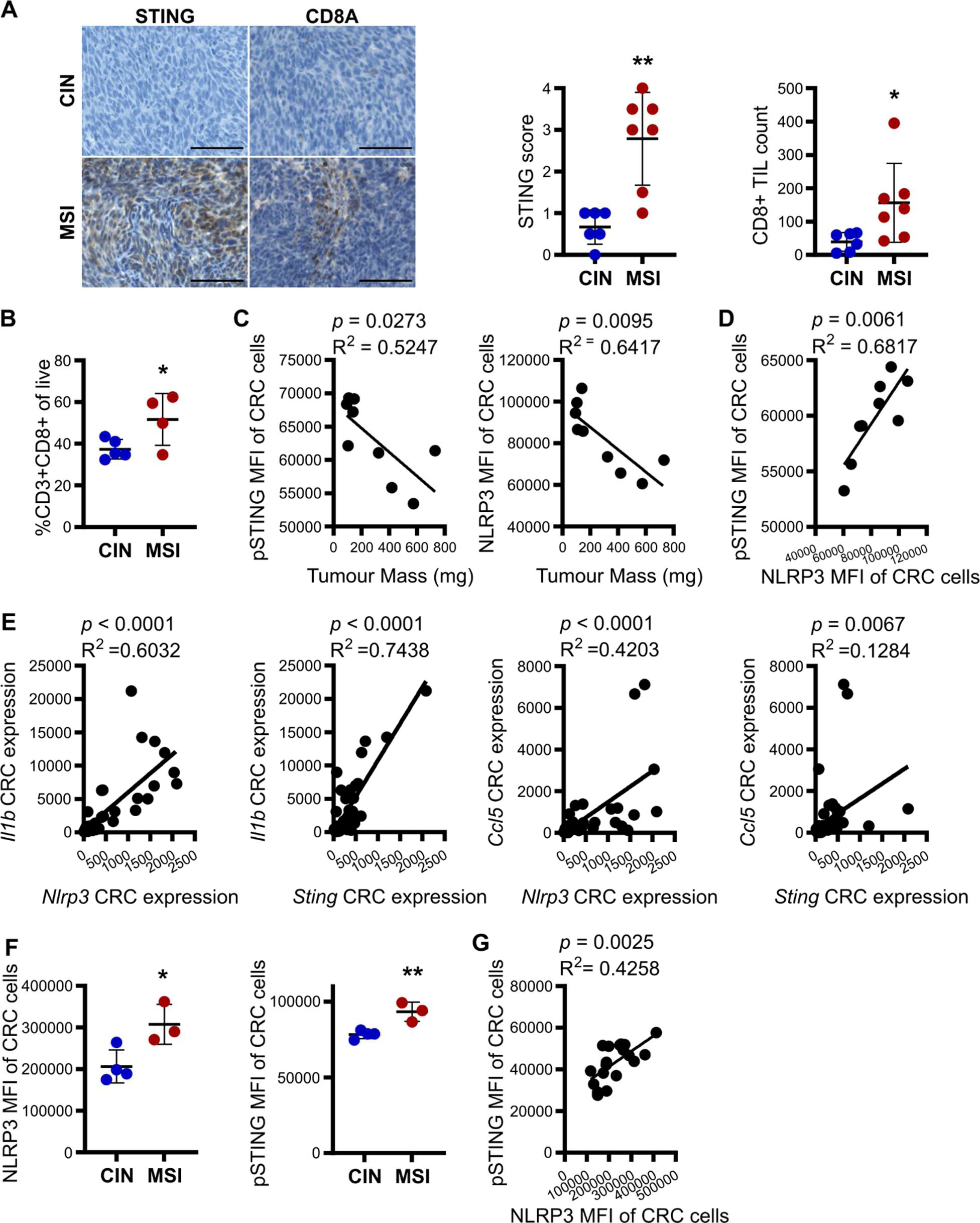
*NLRP3* and *STING* are co-expressed and associated with *CD8A* expression in a murine MSI and CIN CRC model. **(A)** STING and CD8A expression in subcutaneously grown MSI and CIN MC38 CRC tumors as evaluated by IHC. Scale bar = 100 µm. **(B)** CD8+ T cell infiltration into tumors as evaluated by flow cytometry. **(C-D)** Correlation of phospho-STING (pSTING) and NLRP3 expression in the CRC cells as measured by flow cytometry with tumor size and with each other in all mice. **(E)** Correlation between the expression of indicated genes in CRC cells isolated from subcutaneously grown tumors. CRC cells were isolated and qPCR was performed on RNA extracted from these cells. **(F)** Expression of pSTING and NLRP3 in CRC cells of MSI and CIN MC38 CRC tumors grown orthotopically in the colons of immunocompetent mice. Tumors were harvested and analyzed by flow cytometry. **(G)** Correlation of pSTING and NLRP3 expression in the CRC cells of orthotopically grown CRCs. All panels show representative data from N ≥ 3 experiments. Panels A, B, F: \**p* < 0.05, \*\**p* < 0.01. Panels C-E, G: Linear regression (C.I. = 95%) with the Pearson correlation R^2^ and *p*-values reported on each graph.

The subcutaneous tumor model allowed us to perform an initial evaluation of the immune consequences of *Mlh1* loss, but lacked the immune suppressive environment of the mucosal immune system and exposure to the intestinal microbiota known to regulate intestinal immune function and, especially, PRR activation.^10,17^ We thus moved to an orthotopic model where the MSI and CIN MC38 CRC cells are endoscopically injected into the colonic wall of immunocompetent mice.^16^ MSI CRCs in this model had a higher percentage of NLRP3 and STING expressing tumor cells, suggesting that MSI CRCs are more sensitive to microbial signals that activate these pathways (**Fig. 2F**). Regardless of tumor type, expression of pSTING and NLRP3 in the CRC cells was highly correlated (**Fig. 2G**). Collectively, these results indicate that activation of the NLRP3 inflammasome may be an important contributor to cGAS/STING-mediated antitumor immunity in CRC and may contribute to TIL activation in MSI CRCs.

### NLRP3 and STING cooperatively regulate each other’s signaling and expression

Evidence indicates that activation of the cGAS/STING pathway can upregulate transcript levels for the NLRP3 inflammasome substrate pro-*IL1B*.^18^ In order to further investigate the connection between these two signaling pathways, we stimulated MSI and CIN MC38 CRCs with either cGAMP or ATP to activate STING and NLRP3, respectively, or a combination of cGAMP and ATP. As shown in **Fig. 3**, dual stimulation clearly upregulated mediators of both signaling pathways more than the pathway-specific ligands alone. In particular, addition of ATP to cGAMP induced stronger activation of STING itself as well as of TBK1 and NFKB (**Fig. 3A** and **B**) as well as increased NLRP3 inflammasome assembly as measured by ASC-NEK7 speck formation (**Fig. 3D** and **E**). Furthermore, addition of ATP to cGAMP greatly upregulated secretion of the chemokines CCL5 and CXCL10, which are required for CD8+ T cell recruitment into MSI CRCs (**Fig. 3C**). Dual activation could be reversed by addition of the STING inhibitor H151 (**Fig. 3B** and **C**) or the NLRP3 inhibitor MCC950 (**Fig. 3D**) but was not enhanced by addition of the NLRP3 inflammasome product IL18 (**Fig. 3C**), indicating that interaction between these pathways relies on initial intracellular mechanisms and not on later downstream signaling induced by gene expression changes. Notably in most cases, dual stimulation led to equal activation of the cGAS/STING and NLRP3 pathways in both MSI and CIN CRC cells. This confirms our recent finding that there are no fundamental defects in these signaling pathways within CIN CRC cells and that they are equally sensitive to external ligands despite having lower baseline levels due to their lower production of endogenous ligands than MSI CRC cells.^16^

**Figure 3.**
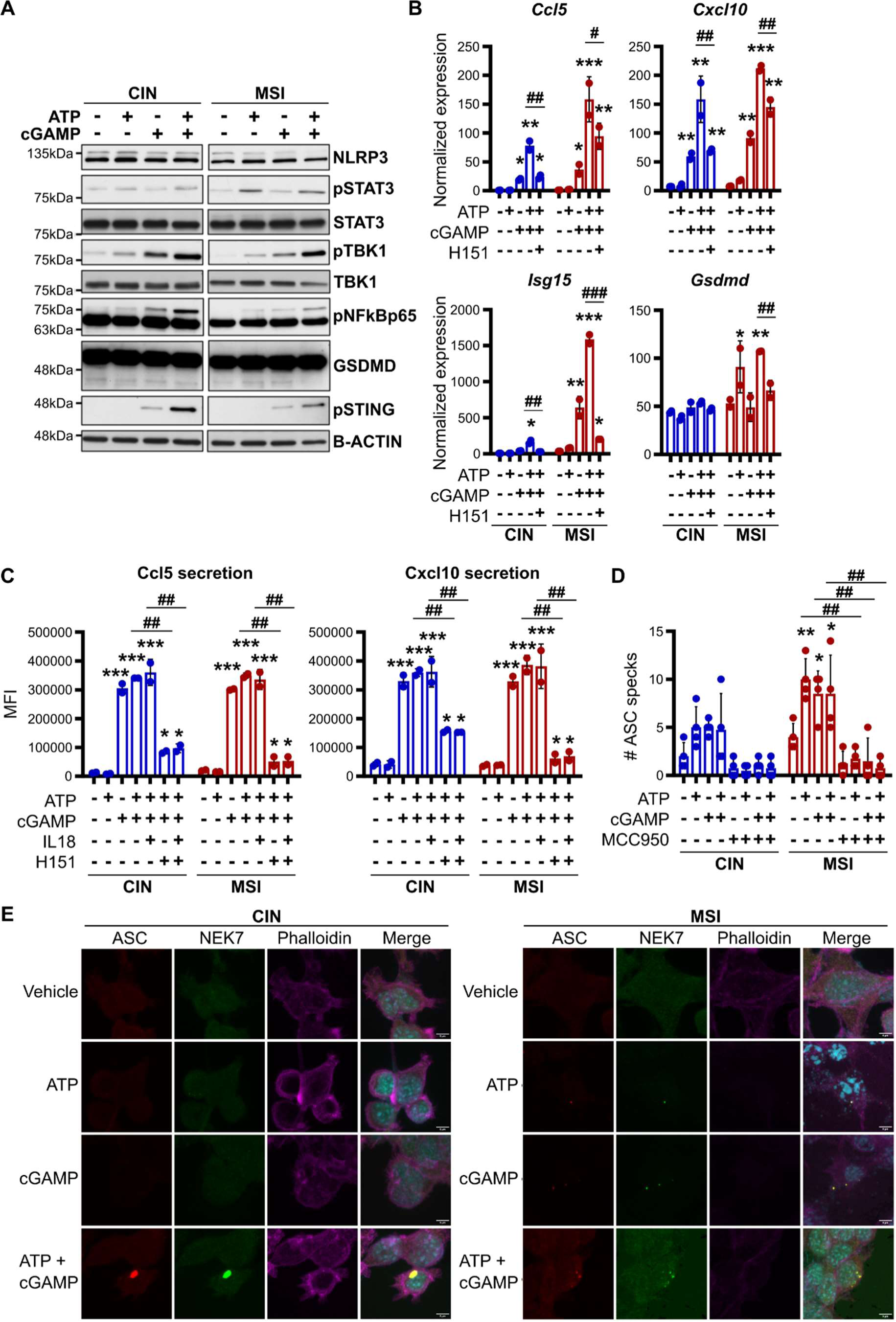
The cGAS/STING and NLRP3 pathways cooperatively regulate each other. **(A-C)** MSI or CIN MC38 CRC cells were stimulated for 24 h with 9 µg/ml 2’-3’-cGAMP and/or 2 mM ATP. Activation of signaling pathway mediators was assessed by Western blotting (A) and expression of downstream products was assessed by qPCR (B) or flow cytometric bead assay of cell supernatants (C). Where indicated, cells were first pretreated for 1 h with 2 µM H151, 10 µM MCC950 or 0.1 µg/ml IL18. **(D-E)** MSI and CIN CRC cells were stimulated as indicated for 4 h and then evaluated by confocal microscopy for colocalization of ASC and NEK7 in ASC specks. Quantification is shown in (D) and representative images in (E). Scale bar = 5 µm. All panels show representative data from N ≥ 3 experiments each with ≥ 2 biological replicates. For all panels, relative to the untreated control: \**p* ≤ 0.05, \*\**p* ≤ 0.01, \*\*\**p* ≤ 0.001. For all panels, between indicated samples: ^#^*p* ≤ 0.05, ^##^*p* ≤ 0.01, ^###^*p* ≤ 0.001.

Despite the clear cooperative effect we observed between cGAS/STING and NLRP3, ATP stimulation alone did not appear to induce cGAS/STING pathway activation (**Fig. 3A-C**). It thus appears that STING and NLRP3 activation work together to enhance STING activity, rather than STING simply increasing expression of NLRP3 or its substrates, which then act independently. To directly test this, we knocked-down *Sting* in MSI MC38 CRC cells (MSI*^StingKD^*). This abrogated not only activation of the cGAS/STING pathway (**Fig. 4A** and **B**) but also decreased expression of the NLRP3 products pro-*IL18* (**Fig. 4B**) and cleaved caspase-1 (**Fig. 4A**). Furthermore, when the MSI*^StingKD^* tumors were orthotopically implanted into the colons of immunocompetent mice, the tumor cells expressed lower amounts of NLRP3 and recruited fewer activated CD8+ T cells than did the control tumors (**Fig. 4C**). Our data thus suggest a dual mechanism where cGAS/STING activation regulates mediators of NLRP3 signaling that then act in a positive feedback loop to directly increase STING pathway activation.

**Figure 4.**
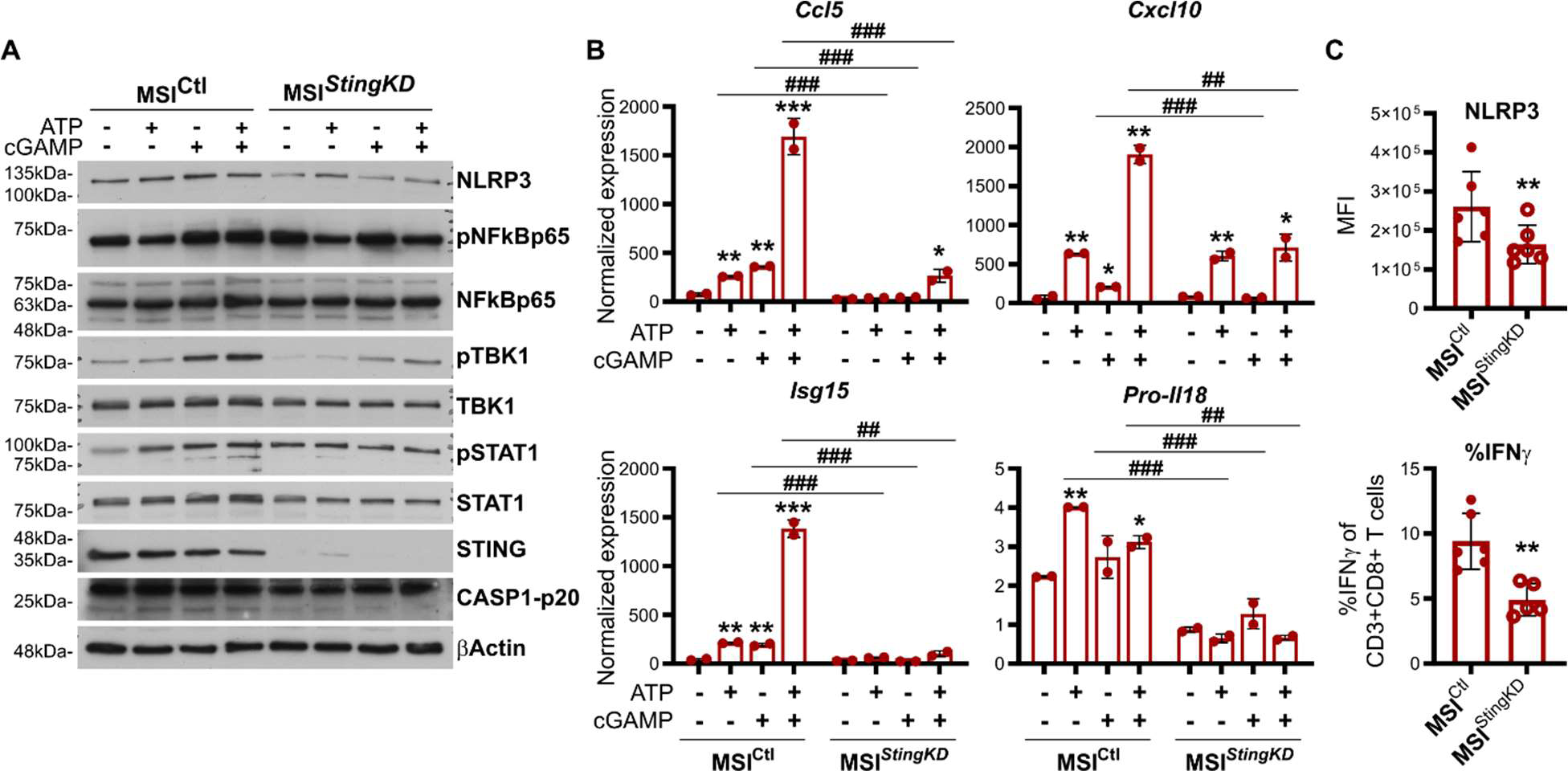
NLRP3 signaling requires activation of the cGAS/STING pathway. *Sting* expression was stably knocked down in the MSI MC38 CRCs to generate MSI*^StingKD^* cells. **(A-B)** Cells were stimulated for 24 h with 9 µg/ml 2’-3’-cGAMP and/or 2 mM ATP and activation of signaling pathways (A) was evaluated by Western blotting while expression of downstream products (B) was evaluated by qPCR (B). **(C)** CRC cells were grown orthotopically in the colons of immunocompetent mice and then tumors were harvested and evaluated by flow cytometry for NLRP3 expression or activation of infiltrating CD8+ T cells. All panels show representative data from N ≥ 3 experiments each with ≥ 2 biological replicates. For all panels, relative to the untreated control: \**p* ≤ 0.05, \*\**p* ≤ 0.01, \*\*\**p* ≤ 0.001. For all panels, between indicated samples: ^##^*p* ≤ 0.01, ^###^*p* ≤ 0.001.

Given that cells contain many additional PRRs that could potentially influence STING signaling, we tested whether activation of two additional PRR pathways could replicate the effect we observed with NLPR3. Stimulation of CRC cells with poly(dA:dT), an agonist of several PRRs including the AIM2 inflammasome, but not of NLRP3, significantly increased expression of the STING pathway products *Ccl5*, *Cxcl10* and *Isg15* in addition to the inflammasome target pro-*Il18* (Supplementary Fig. 2A).^19^ However, there was no cooperativity between AIM2 and cGAS/STING since addition of cGAMP did not increase activation beyond that of poly(dA:dT) alone. In contrast, stimulating our CRC cells with both cGAMP and flagellin, a TLR5 agonist, did not increase expression of *Ccl5*, *Cxcl10*, *Isg15* and pro-*Il18* more than stimulating with cGAMP alone (Supplementary Fig. 2B).^20^ It thus appears that specific PRRs, especially inflammasomes, can promote cGAS/STING pathway signaling alone but that cooperativity with cGAS/STING signaling is not a general feature of all PRRs. Since the main outcome of canonical NLRP3 inflammasome stimulation is activation of caspase-1, we tested whether interaction between the NLRP3 and the cGAS/STING pathways relies on caspase-1 activation (Supplementary Fig. 2C). However, addition of the caspase-1 inhibitor YVAD to the dual stimulation of CRC cells did not decrease cGAS/STING activation by NLRP3, indicating that the interaction is caspase-1-independent.^21,22^ Given that most effects of NLRP3 depend on caspase-1 activation, our data indicate that there is a great deal of specificity in the cooperativity between cGAS/STING and NLRP3 and that dual stimulation of these pathways can produce very specific immune effects rather than promoting generalized inflammation.

### NLRP3 and cGAS/STING activation in CRC cells cooperatively regulate activation of CD8+ T cells

NLRP3 inflammasome activation is typically associated with tumor-promoting inflammation. To verify that activating NLRP3 in CRC cells could specifically potentiate cGAS/STING-mediated activation of anti-tumor CD8+ T cells, we directly tested if simultaneously stimulating these two pathways in MSI and CIN CRC cells could activate CD8+ T cells. We stably transduced *Mlh1^-/-^*and *Kras^Mut^* MC38 CRC cell lines with the model tumor neoantigen ovalbumin (OVA) to generated MSI^OVA^ and CIN^OVA^ cells, respectively. We then pre-treated these cells with ATP and/or cGAMP for 24 h, removed the treatment, and then cocultured the CRCs with OVA-specific OT-I CD8+ T cells. Both MSI^OVA^ and CIN^OVA^ CRC cells treated with the NLRP3 ligand ATP alone activated significantly more CD8+ T cells than did the vehicle control cells (**Fig. 5A**). Similarly, NLRP3 activation alone potentiated CD8+ T cell-mediated tumor cell killing. Thus, NLRP3 stimulation alone can induce a productive antitumor immune response in both MSI and CIN CRC cells, although the effect was strongest in MSI CRCs. Dual stimulation of the CRC cells with cGAMP and ATP even further increased activation of CD8+ T cells in response to the tumor cells (**Fig. 5A**). To confirm these data and determine the extent to which this effect was dependent on the ability of NLRP3 to directly activate the cGAS/STING pathway, we stimulated MSI*^StingKD^* CRCs cells with ATP, cGAMP or both. NLRP3 activation failed to enhance CD8+ T cell activation by these STING-deficient cells despite activating CD8+ T cell activation by the control cells (**Fig. 5B**). CD8+ T cell activation by NLRP3 thus appears to require STING. This is consistent with our observation that orthotopically injected MSI*^StingKD^* CRC cells express lower NLRP3 and attract fewer activated IFNγ-expressing CD8+ T cells than control MSI CRC cells (**Fig. 4C**). Collectively, these data show that dual activation of both STING and NLRP3 cooperatively induces antitumor immunity in MSI CRC cells and suggests that one way to maximize CD8+ T cell activation in CIN CRC patients would be to treat them with therapeutics that stimulate both pathways.

**Figure 5.**
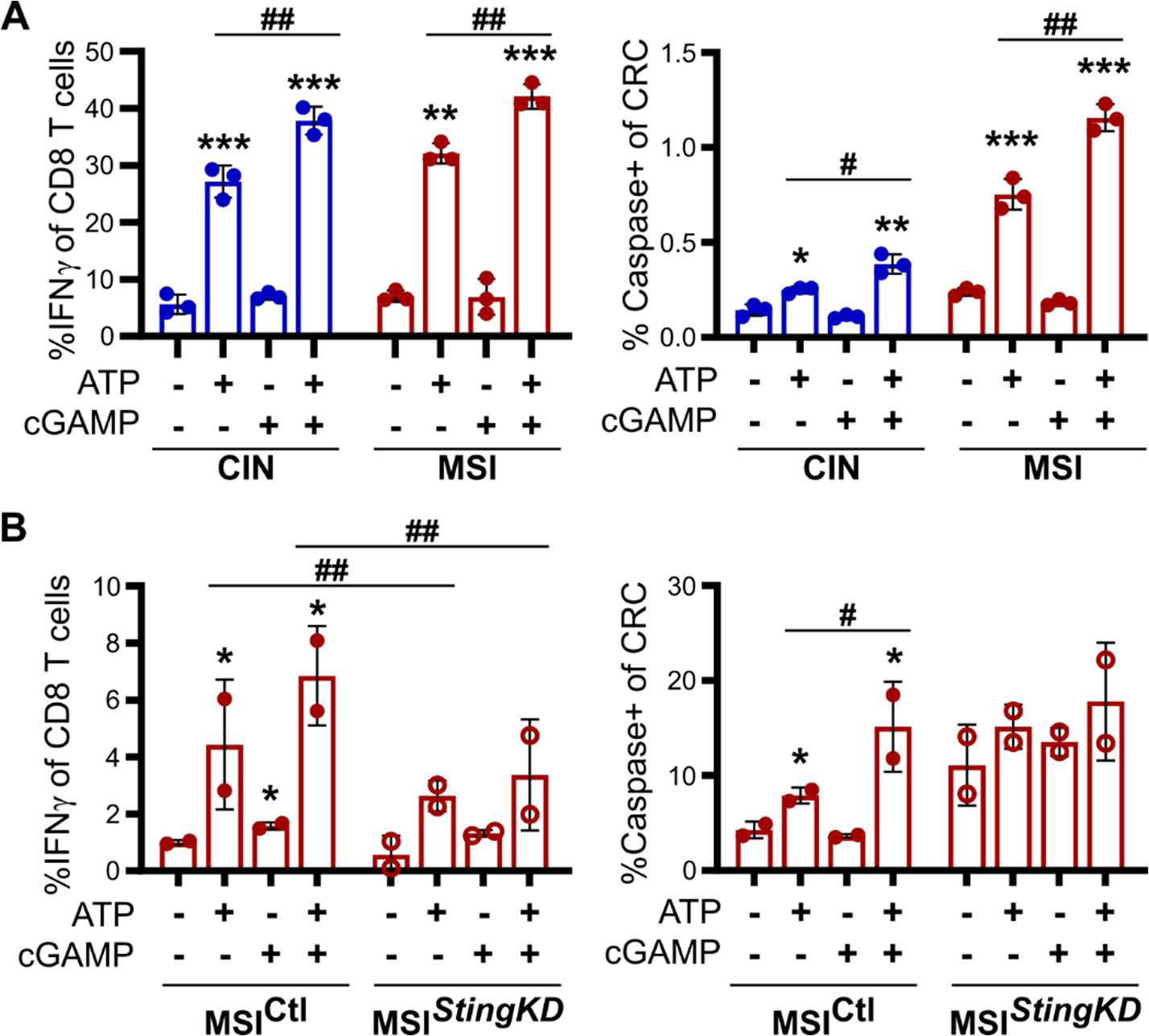
cGAS/STING and NLRP3 activation in CRC cells cooperatively regulate activation of CD8+ T cells. OVA-expressing MSI and CIN MC38 CRC cells were stimulated for 24 h with 9 µg/ml 2’-3’-cGAMP and/or 2 mM ATP. The cells were then washed and cocultured for an additional 48 h with OT-I CD8+ T cells. **(A-B, left panels)** Activation of CD8+ T cells was measured by intracellular flow cytometry. **(A-B, right panels)** T cell-mediated killing of the CRC cells was measured by caspase 3/7 cleavage. All panels show representative data from N = 3 experiments each with 2-3 biological replicates. For all panels, relative to the untreated control: \**p* ≤ 0.05, \*\**p* ≤ 0.01, \*\*\**p* ≤ 0.001. For all panels, between indicated samples: ^#^*p* ≤ 0.05, ^##^*p* ≤ 0.01.

### Activating cGAS/STING and NLRP3 in CRC increases antitumor immunity in orthotopic CRC

Activation of STING signaling is well known to promote CD8+ T cell-mediated antitumor immunity in vivo but little has been reported to this effect for NLRP3.^13,16,23^ To test this, we orthotopically injected MSI and CIN MC38 CRCs into the colons of immunocompetent mice and then treated them repeatedly with the NLRP3 inhibitor MCC950 (**Fig. 6A**).^14,24^ This significantly decreased the activation of CD8+ T cells in MSI CRCs and the draining mesenteric lymph nodes (MLNs) (**Fig. 6B**). This effect was particularly strong for the MSI CRCs, consistent with our finding of greater baseline activation of this pathway in MSI CRCs. This documents a previously unrecognized role for NLRP3 in CRC antitumor immunity. Since cGAS/STING signaling was already known to promote CD8+ T cell-mediated tumor immunity, we specifically sought to test whether doing so therapeutically could enhance the performance of anti-PD1 immune checkpoint inhibitor (ICI) therapy in tumors growing in a complex environment such as the colon. Mice with orthotopically growing MSI and CIN CRCs were injected at regular intervals with anti-PD1 ICI or an isotype control in combination the STING agonist ADU-S100 (**Fig. 6C**).^25,26^ Upon harvesting the tumors, we noted a small increase in CD8+ T cell activation in the MSI tumors when mice received both anti-PD1 and S100A (**Fig. 6D**, Supplementary Fig. 3A). However, when we examined the MLNs, which are the predominant site of T cell priming in the intestine, there was significantly higher CD8+ T cell activation in both the MSI and CIN CRCs in mice receiving the dual treatment (**Fig. 6E**). We also noted a decrease in MHCII but not MHCI expression on MLN DCs, which is consistent with their polarization towards a CD8+ T cell-priming phenotype (Supplementary Fig. 3B). In addition, we noted much higher induction of the Cxcl10 receptor, Cxcr3, on CD8+ T cells in the MLNs and spleen of both MSI and CIN CRCs treated with dual stimulation (**Fig. 6F,G**). This is significant because we have previously shown that CXCL10-induced trafficking is essential for recruitment of CD8+ T cells into MSI CRCs and could thus explain why we saw more activated T cells in the MSI but not the CIN tumors. It also implies that the strategy of dual stimulation of separate pathways can promote antitumor immunity in otherwise refractory type cancers.

**Figure 6.**
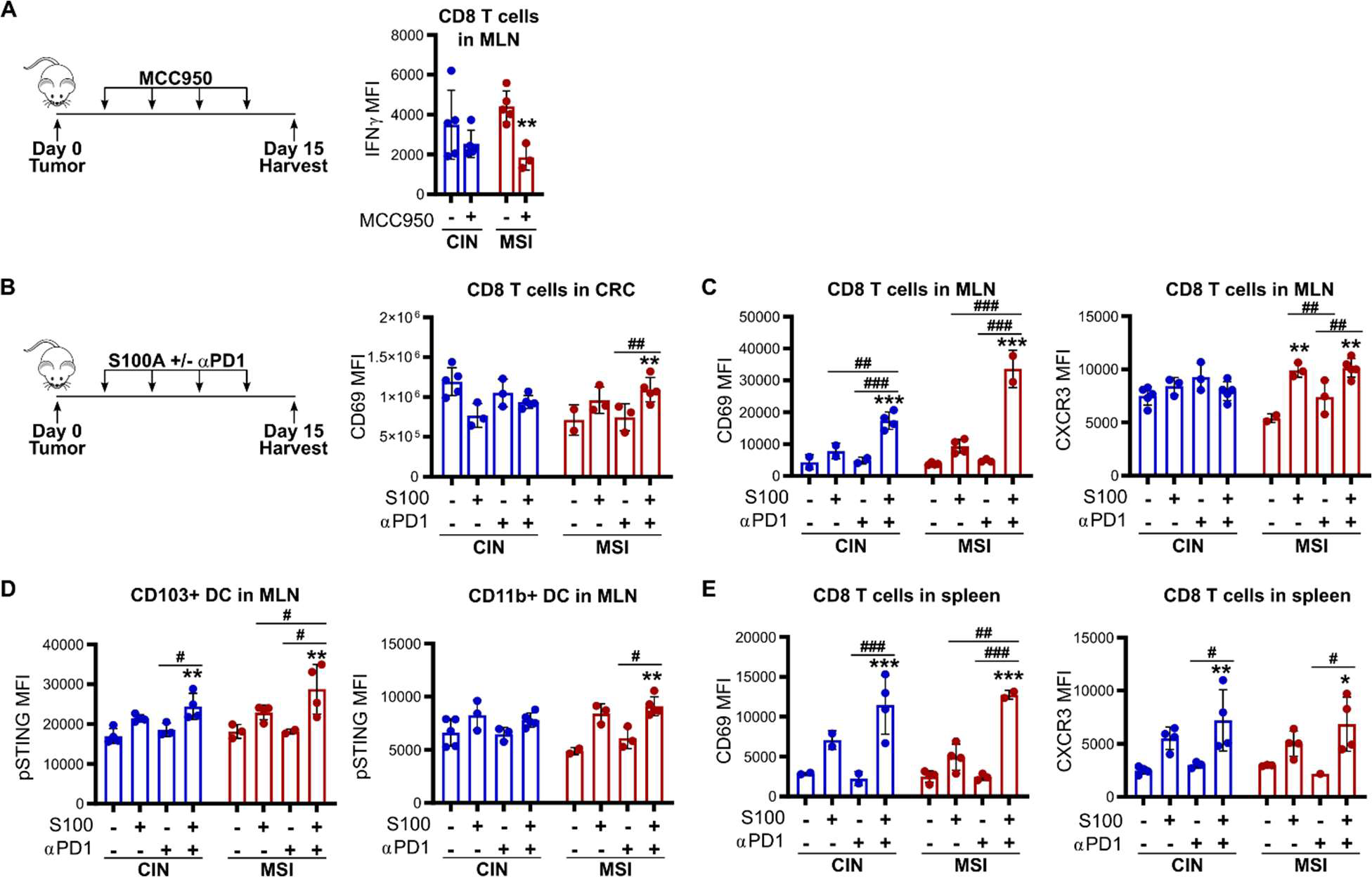
Activating cGAS/STING and NLRP3 in CRC increases antitumor immunity in orthotopic CRC. **(A)** MC38 CRC cells were orthotopically implanted into the colons of immunocompetent mice. Mice were then treated every 3 days with 20 mg/kg MCC950 via IP injection before harvesting tumors, MLNs and spleen for flow cytometric analysis. **(B-E)** MC38 CRC cells were orthotopically implanted into the colons of immunocompetent mice. Mice were then treated every 3 days with 1 mg/kg ADU-S100 and 200 µg of anti-PD1 or isotype via IP injection before harvesting tumors, MLNs and spleen for flow cytometric analysis. (B) CD8+ T cell activation in the CRCs. (C) CD8+ T cell activation in the MLNs. (D) pSTING expression in CD103+ DCs in the MLNs. (E) CD8+ T cell activation in the spleen. All panels show representative data from N = 3 experiments each with 3-5 mice per experiment. For all panels, relative to the untreated control: \**p* ≤ 0.05, \*\**p* ≤ 0.01, \*\*\**p* ≤ 0.001. For all panels, between indicated samples: ^#^*p* ≤ 0.05, ^##^*p* ≤ 0.01, ^###^*p* ≤ 0.001.

### NLRP3 and cGAS/STING cooperatively regulate antitumor immunity in primary mouse and human CRCs

Although cell lines are extremely valuable tools, we sought to confirm that cooperativity between NLRP3 and cGAS/STING activation was not an artifact of the MC38 CRC cell line but a bona fide biological mechanism. To do this, we established primary organoids from CRCs induced in wild type mice by repeated administration of azoxymethane (AOM).^16,27,28^ We created an MSI variant of these organoids *Mlh^KD^*) by stably knocking down *Mlh1*. As seen with the MC38 CRC cells, dual stimulation increased expression of *Ccl5* and *Cxcl10* more than either agonist alone, especially for the *Mlh1^KD^* cells (**Fig. 7A**). In addition, we observed stronger activation of OT-I CD8+ T cells by the MSI organoids than the control CIN-like organoids (**Fig. 7B**) which was enhanced by activation of either NLRP3 or STING alone and even more strongly enhanced by dual stimulation of both pathways (**Fig. 7C**).

**Figure 7.**
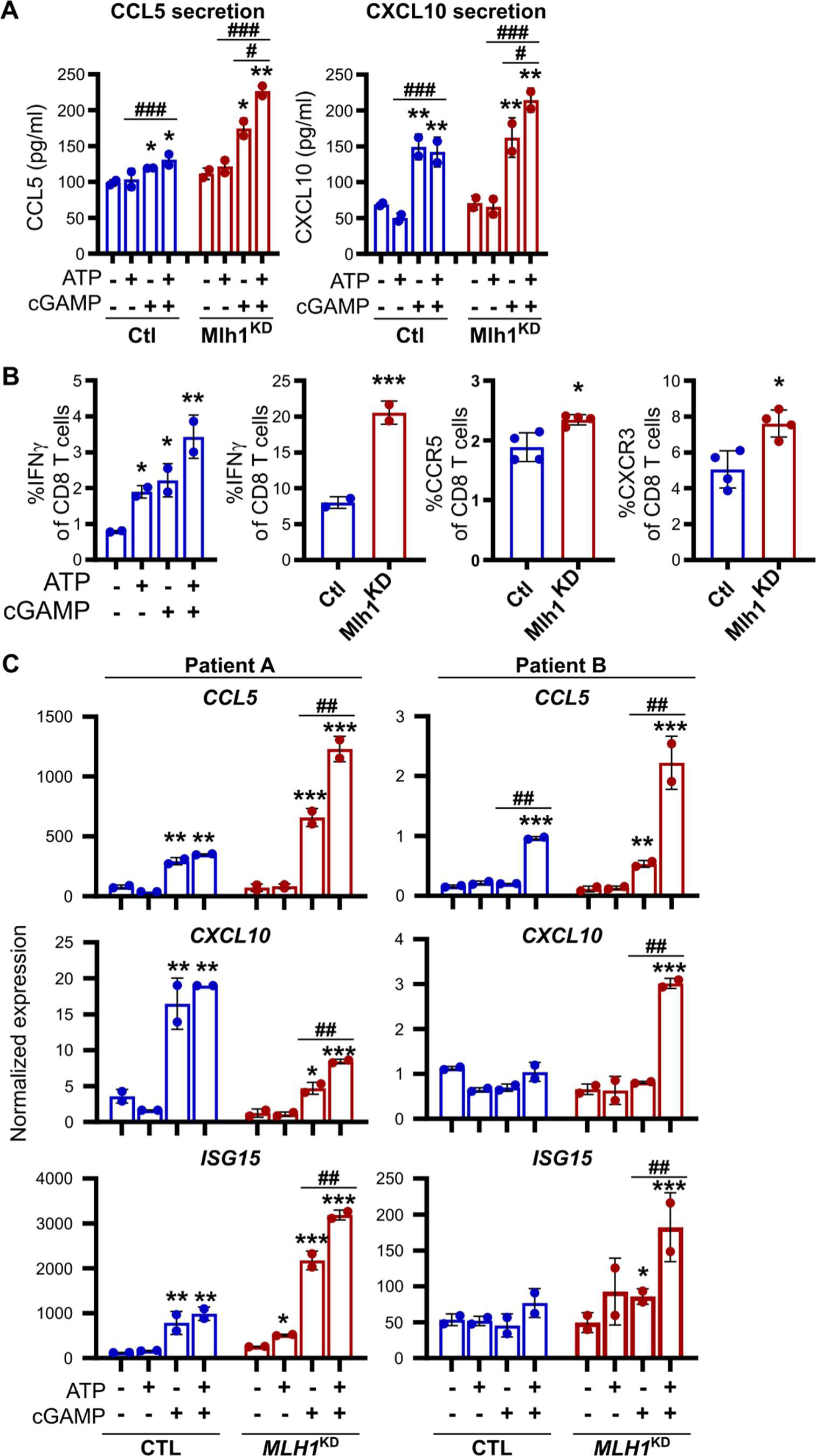
NLRP3 and cGAS/STING cooperatively regulate antitumor immunity in primary mouse and human CRC cells. **(A,B)** Primary organoids were generated from CRC tumors induced in mice by repeated injection of AOM. *Mlh1* was then stably knocked down to create an MSI variant (Mlh1^KD^). (A) Organoids were stimulated for 24 h with 9 µg/ml 2’-3’-cGAMP and/or 2 mM ATP before their supernatant was harvested and secreted cytokines were assessed by flow cytometric bead array. (B) Organoids were treated as in A and then washed before pulsing with SIINFEKL and coculturing with OT-I CD8+ T cells for 48 h. T cell activation was assessed by flow cytometry. **(C)** Primary organoids were generated from the resected tumors of two different CRC patients. *MLH1* was then stably knocked down in each to create an MSI variant (MLH1^KD^). Organoids were stimulated for 24 h with 9 µg/ml 2’-3’-cGAMP and/or 2 mM ATP. RNA was then collected and gene expression was evaluated by qPCR. All panels show representative data from N ≥ 3 experiments each with 2-4 biological replicates. For all panels, relative to the untreated control: \**p* ≤ 0.05, \*\**p* ≤ 0.01, \*\*\**p* ≤ 0.001. For all panels, between indicated samples: ^#^*p* ≤ 0.05, ^##^*p* ≤ 0.01, ^###^*p* ≤ 0.001.

To further extend the significance of NLRP3 and cGAS/STING cooperativity, we generated primary organoids from human CRCs and, for each patient’s samples, created an MSI variant by knocking-down *MLH1* expression (*MLH1^KD^*). Stimulation of either STING or NLRP3 upregulated expression of STING-associated chemokines *CCL5*, *CXCL10* and *ISG15* (**Fig. 7D**). Expression of each of these was even further enhanced by dual stimulation of STING and NLRP3 in both the MSI and CIN organoid variants from each patient (**Fig. 7D**). These findings confirm that dual stimulation of NLRP3 and cGAS/STING pathways is an attractive therapeutic strategy for increasing the ability of CIN CRC cells to activate CD8+ T cell-mediated antitumor immunity.

## Discussion

Most CRCs are poor stimulators of antitumor immunity and are resistant to currently available immunotherapies. We show here that a promising therapeutic strategy for enhancing CD8+ T cell activation in CRC is dual stimulation of the cGAS/STING and NLRP3 pathways which act cooperatively to enhance CD8+ T cell immunity. Specifically, cGAS/STING signaling regulates the expression of NLRP3 pathway mediators and NLRP3 directly activates cGAS/STING signaling, thereby setting up a positive feedback loop that boosts secretion of the T cell recruiting chemokines CCL5 and CXCL10 as well as increases activation of CD8+ T cells following their infiltration into the tumors. While we show that dual activation of these pathways is strongest in the immunogenic MSI CRC subtype, CIN CRCs also significantly increased T cell activation in response to treatment.

Inflammation is very much a dual-edge sword in cancer.^29,30^ While high grade and chronic inflammation both predispose to cancer and accelerate its progression, low level acute inflammation can potentiate the activation of tumor-targeting cytotoxic T cells by providing necessary costimulatory signals. While formation of the NLRP3 inflammasome is typically associated with chronic inflammation, our findings illustrate that it can also enhance CD8+ T cell activation by feeding into the cGAS/STING pathway and increasing production of IFN-stimulated genes. This provides important insight into how crosstalk between innate signaling pathways can fine tune the overall tumor immune microenvironment. It is possible that baseline activation of other PRRs such as cGAS/STING determines the consequence of NLRP3 signaling. This might explain why MSI CRCs, which have higher endogenous cGAS/STING activation than CIN CRCs, responded more strongly to dual stimulation of NLRP3. Given the complex composition of the infectious agents that have largely driven evolution of the PRR system, such integration would be a natural way to fine tune the immune response towards a particular invading pathogen. This could explain how a potentially detrimental inflammatory response could instead produce successful CD8+ T cell activation in some circumstances. Exploiting existing crosstalk between these signaling pathways could thus prove a powerful way to increase the immunogenicity of CIN CRCs while minimizing induction of inflammatory adverse events.

The ability of cGAS/STING signaling to activate the NLRP3 inflammasome is well documented and seems to involve both upregulation of NLRP3 pathway mediators as well as the induction of potassium efflux that triggers inflammasome assembly.^31–33^ This is consistent with our findings of correlation between cGAS/STING activation and NLRP3 expression in the CRC cells in addition to the increase in ASC specks we identified upon dual stimulation. In contrast, the mechanism by which NLRP3 enhances cGAS/STING signaling is not yet clear and contradicts several previous studies that have found NLRP3 activation to repress cGAS/STING signaling.^34,35^ One possibility is that activation of the NLRP3 inflammasome could lead to caspase-1-mediated DNA damage, including mitochondrial DNA oxidation, that results in release of the damaged DNA into the cytosol where cGAS can detect it.^36–38^ However, our data indicate that the caspase-1 inflammasome is not required for amplification of cGAS/STING signaling by NLRP3 and that it instead proceeds via an inflammasome-independent function of NLRP3.^39,40^ These largely involve generating reactive oxygen species (ROS) in the mitochondria and inducing mitophagy and would be expected to generate damaged DNA that could be sensed by cGAS.^33,41–43^ In addition, NLRP3-mediated cGAS/STING activation could result from its role as a stress sensor, especially in MSI CRCs that experience high levels of genotoxic or ER stress induced by excessive point mutations and aberrant proteins.^32,44^ Alternatively, NLRP3 could be activated by signaling from the high levels of immune cells that infiltrate MSI CRCs or by specific microbial taxa that have been found to be enriched in MSI CRCs.^45–48^

Further research is clearly needed to uncover the detailed mechanism by which innate immune sensors such NLRP3 can contribute to regulation of productive antitumor immunity while minimizing their contribution to tumor-promoting inflammation. Our current work shows how two such pathways interact to boost CD8+ T cell activation and that dual activation of these pathways can render otherwise immunologically refractory tumors susceptible to T cell-mediated killing. Developing therapies that exploit the crosstalk between these pathways is thus a promising strategy to maximize response rates and optimize the performance of existing and future cancer immunotherapies.

## Materials and Methods

### Cell lines

MC38 murine CRC cells were purchased from Kerafast and maintained in high glucose DMEM supplemented with 10% FBS, 1% penicillin-streptomycin, and 1% HEPES at 37°C with 5% CO_2_. MSI and CIN variants of the cells were generated using CRISPR-Cas9 editing to delete *Mlh1* or mutate *Kras*, respectively, as described previously.^16,49^ *Sting* knockdown in the MSI MC38 CRC cells was generated as described previously using shRNA and the pLKO.1 system.^16,50^ Ovalbumin (OVA) expressing MSI and CIN MC38 CRC cells were made by transfection with the pCI-neo-cOVA plasmid (Addgene #25097) and selection with 200 µg/ml G418.^51^

### Cell stimulations

MC38 CRC cells were seeded into plates 24 h before treatments as indicated in the figure legends. The following treatments were used: 9 µg/ml 2’,3’-cGAMP, 2 mM ATP (SigmaAldrich), or 0.1 µg/ml IL18 (BioLegend). cGAMP was delivered to the cells encapsulated in Lipofectamine 2000 (ThermoFisher). In some cases, cells were first pre-treated for 1 h with following inhibitors (2 µM H-151, 10 µM MCC950 (InvivoGen)) before addition of the treatments. After the indicated incubation time, cells supernatant or cells were harvested as indicated below.

### Immunofluorescence staining

Cells were seeded onto coverslips coated with CellTak and allowed to adhere overnight at 37°C. Cells were fixed with methanol and blocked (5% normal serum, 0.3% Triton X-100 in PBS). Cells were stained with the indicated primary antibodies for 1 h at room temperature. After washing, cells were stained with secondary antibodies (ThermoFisher) for 1h at room temperature followed by phalloidin Alexa^546^ (ThermoFisher) for 20 min. Nuclei were stained with DAPI and then coverslips were mounted on slides. Cells were imaged on a Leica Falcon SP8 STED System. Post image processing was performed using Image J.^52^

### Western blotting

Protein was isolated in lysis buffer (50 mM Tris-HCl, 150 mM NaCl, 50 mM sodium pyrophosphate, 1 mM EDTA, 0.5% NP40, 1% Triton X-100) containing 1mM sodium orthovanadate, and 1x protease inhibitor (SigmaAldrich). Protein was quantified using a BCA assay (ThermoFisher) and 5 or 10 µg of protein was loaded on SDS-PAGE gels and transferred to nitrocellulose membranes. See Supplementary Table 1 for all antibodies. Bands were visualized using the ECL Prime Western Blotting Detection Reagent (Cytiva).

### RNA and qPCR

RNA was extracted using Trizol and reverse transcribed using the High-Capacity cDNA Reverse Transcription Kit (ThermoFisher). qPCR was performed with the primers in Supplementary Table 1 using the POWRUP SYBR Master Mix (ThermoFisher). qPCR was performed on the QuantStudio6 real-time PCR system (Applied Biosystems).

### Mouse CRC experiments

C57BL/6 wildtype mice originally purchased from Charles River were bred and maintained in the Cross Cancer Institute vivarium. OTI mice were purchased from The Jackson Laboratory. Mixed groups of male and female littermates between the age of 6-20 weeks old were used for experiments. All animal work was approved by the Cross Cancer Institute’s Animal Care Committee.

Subcutaneous CRC experiments were performed by injecting 5×10^5^ MSI or CIN MC38 CRC cells in 100 µl PBS into the hind flank. Tumors were harvested after 2-3 weeks. Tumors were dissociated in an enzyme cocktail of RPMI containing 0.5 mg/ml collagenase IV (SigmaAldrich), 10 µg/ml DNaseI (SigmanAldrich), 10% FBS, 1% penicillin-streptomycin and 1% HEPES buffer for 30 minutes at 37°C in a shaking incubator at 200 rpm.^16,27^ Fragments were rigorously pipetted to dissociate, filtered through a 100 µm cell strainer and washed. Cancer and immune cells in the tumors were separated using a 40%/80% percoll gradient (Cytiva), at 500 g for 30 min at room temperature. The top layer (tumor cells) and the interface (immune cells) were collected separately, washed, and processed for RNA isolation as below.

Orthotopic CRC experiments were performed by injecting 1.5×10^5^ MC38 CRC cells in 50 µl PBS into the wall of the descending colon using a flexible needle (Hamilton) inserted through the working channel of a Wolfe endoscope and visualized via the ColoView imaging system (Storz).^16,53^ In some experiments, mice were treated intraperitoneally every 3 days with either 1 mg/kg ADU-S100 and 200 µg anti-PD1/isotype (BioXcell) or with 20 mg/kg MCC950. Orthotopic tumors, MLNs and spleen were harvested after 14-21 days and dissociated as above before flow staining.

### Mouse and human CRC-derived organoids

Murine organoids were generated from colorectal tumors induced in wildtype C57BL/6 mice by repeated doses of azoxymethane (10 weekly doses of 10 mg/kg azoxymethane) as previously described and cultured as below.^16,28,54^

Human CRC organoids were generated as described previously from resected tumors.^16,54^ Briefly, tumors were dissociated for 1 h in DMEM with 2.5% FBS, 75 U/ml collagenase XI (SigmaAldrich), 125 µg/ml dispase II (SigmaAldrich). Following filtration, cells were plated at 500-1000 per well in growth factor reduced Matrigel (Corning) and cultured in basal crypt media (Advanced DMEM/F12containing 10% FBS, 2 mM glutamine, 10 mM HEPES, 1 mM N-acetylcystein, 1X N2 supplement, 1X B27 supplement, 10 mM nicotinamide, 500 nM A83-01, 10 µM SB202190, 50 ng/ml EGF) (ThermoFisher) mixed 1:1 with conditioned supernatant from L-cells expressing Wnt3a, R-spondin and noggin (ATCC #CRL-3276).^55^ All work with human samples was approved by the Health Research Ethics Board of Alberta Cancer Committee and carried out after obtaining informed patient consent.

Primary MSI variants of the mouse and human organoids were generated using lentiviral transduction with the pLKO.1 system and shRNA sequences in Supplementary Table 2.^16,50,53,56^ Briefly, organoids were pretreated for 4-5 days with 10 mM nicotinamide before being dislodged from the plate by pipetting and treated for 5 min at 37°C with TrypLE Express (Life Technologies). Organoids were mixed with concentrated lentivirus along with 8 µg/ml polybrene and 10 µM Y27632 (SigmaAldrich) and seeded into a 96-well plate. The plate was centrifuged for 60 min at 600g at 32°C and then incubated at 37°C for 6 h. The organoids were then embedded in Matrigel and cultured in media containing 50-100 µg/ml hygromycin to select for successful transduction. Gene knockdown was verified by Western blot.

For stimulations, equal numbers of organoids were plated in Matrigel and cultured for 3 days before being treated as indicated in the figure legends. Organoids were harvested by resuspending in ice cold Cell Recovery Solution (CRS) (Corning) to dissolve the Matrigel, diluted in ice cold PBS, spun down and processed for RNA or protein isolation.

### Flow cytometry and cytokine bead arrays

Flow cytometry staining was performed using the antibodies in Supplementary Table 1, the Zombie Aqua viability stain (BioLegend), and the Foxp3 Transcription Factor Staining Buffer Set (eBioscience). Flow cytometry was performed on a CytoFlex S cytometer (Beckman Coulter) with subsequent analysis using FlowJo (BD Biosciences).

Secretion of CCL5 and CXCL10 by MSI and CIN CRCs into cell supernatants was analyzed using a custom LegendPlex cytokine bead array (BioLegend). Data was acquired on a CytoFlex S cytometer and analyzed in FlowJo.

### T cell activation assays

CD8+ T cells were isolated from the spleens and lymph nodes of OTI mice using the EasySep Mouse CD8+ T Cell Isolation Kit (StemCell Technologies). For direct co-culture experiments, OVA-expressing MSI and CIN CRC cells were first treated for 24 h followed by extensive washing. CD8+ T cells were then added at a 5:1 T cell:tumor cell ratio and cultured for 24-48 h. Cytotoxicity was assessed using the CellEvent Caspase-3/7 Green Detection Reagent (ThermoFisher) at 1.0 µM and gating on CD45-negative cells. To assess T cell activation by STING knockdown or scramble controls, CRC cells were treated for 24 h and then pulsed with 1 mg/ml of the SIINFEKL peptide for 30 min before being washed and cultured with OTI CD8+ T cells as above.

### Data from the Cancer Genome Atlas analysis

Human RNA sequencing data and DNA sequencing data (Illumina HiSeq RNASeqV2) from the Colorectal Adenocarcinoma dataset from the TCGA Nature 2012 and TCGA PanCancer Atlas from The Cancer Genome Atlas were downloaded from cBioPortal for Cancer Genomics (https://www.cbioportal.org/).^57–59^ Data was log2 transformed and analyzed using the DESeq2 package in R (v3.0).^60^

### Statistical analysis

Prism (GraphPad Software Inc.) was used for statistical analysis. Comparisons of two unpaired groups was made by two-tailed Student’s t-test for normal data, or Mann-Whitney for non-parametric tests. For three or more groups with two parameters, two-way ANOVA or multiple t-test procedures were used as appropriate, for data with Gaussian distribution. Analysis of responses to multiple stimuli of a single cell type or of cells from a single donor were analyzed using paired tests. Post-hoc analysis to correct for multiple comparisons and detect differences between groups was by the two-stage linear step-up procedure of Benjamin, Krieger and Yekutieli with false discovery rate <0.05. A two-sided probability (p) of alpha error less than 0.05 defined significance.

## Acknowledgements

The authors thank Rose-Marie Cornand, Dan McGinn, Cheryl Santos, Daming Li, Dr. Xuejun Sun, Dr. Anne Galloway and Dr. Aja Rieger as well as the Faculty of Medicine and Dentistry flow cytometry and high throughput cores at the University of Alberta for technical support. This work was supported by funding from the Canadian Institutes of Health Research, the Natural Sciences and Engineering Research Council of Canada, the Canadian Foundation for Innovation, the Cancer Research Society and the University Hospital Foundation (KB).

## Author contributions

C. Mowat conceived of the project, designed and performed experiments, analyzed data and wrote the manuscript. D. Schiller provided human CRC tissue samples. K. Baker conceived of the project, analyzed data, wrote the manuscript and secured funding.

## Notes

**Conflict of Interest:** The authors declare no potential conflicts of interest.

### Competing Interest Statement

The authors have declared no competing interest.

