## Supplementary File for "cGAS/STING and NLRP3 cooperatively activate CD8+ T cell-mediated anti-tumor immunity in colorectal cancer"

**Supplementary Table 1. Antibodies**

| Target | Fluorophore or Secondary | Purpose | Source |
|---|---|---|---|
| b-Actin | Anti-rabbit IgG HRP | Western Blot | Cell Signaling (8457S) |
| GAPDH | Anti-mouse IgG HRP | Western Blot | ThermoFisher (PIMA515738) |
| MLH1 | Anti-rabbit IgG HRP | Western Blot | Abcam (ab92312) |
| Phospho-TBK1 (Ser172) | Anti-rabbit IgG HRP or Alexa 488 | Western Blot, Flow Cytometry | Cell Signaling (5483S) |
| TBK1 | Anti-rabbit IgG HRP | Western Blot | Cell Signaling (3504S) |
| NLRP3 | Anti-rabbit IgG HRP | Western Blot | AbCam (ab263899) |
| Phospho-NFkB p65 (Ser536) | Anti-rabbit IgG HRP | Western Blot | Cell Signaling (3033S) |
| NFkB p65 | Anti-rabbit IgG HRP | Western Blot | Cell Signaling (8242S) |
| Phospho-STAT3 (Tyr705) | Anti-rabbit IgG HRP | Western Blot | Cell Signaling (9131S) |
| STAT3 | Anti-rabbit IgG HRP | Western Blot | Cell Signaling (12640S) |
| Phospho-STAT1 (Tyr701) | Anti-rabbit IgG HRP | Western Blot | Cell Signaling (7649S) |
| STAT1 | Anti-rabbit IgG HRP | Western Blot | Cell Signaling (9172S) |
| Phospho-STING (Ser365) | Anti-rabbit IgG HRP | Western Blot, Flow Cytometry | Cell Signaling (72971S) |
| STING | Anti-rabbit IgG HRP | IHC, Western Blot | Cell Signaling (13647S) |
| Gsdmd | Anti-rat IgG HRP | Western Blot | Genentech (10026) |
| Caspase-1 | Anti-rabbit HRP | Western Blot | AbCam (ab1872) |
| ASC | Alexa488 | IF | Cell Signaling (17507) |
| NEK7 | Anti-rabbit Alexa592 | IF | AbCam (ab133514) |
| CD8A | Anti-rabbit IgG HRP | IHC | Novus (NBP2-29475) |
| CD3 | APCCY7 | Flow Cytometry | Biolegend (100222) |
| CD8A | APC | Flow Cytometry | Biolegend (100712) |
| CD8A | ALEXA-700 | Flow Cytometry | ThermoFisher (56-0081-82) |
| CD4 | PERCPCY5.5 | Flow Cytometry | Biolegend (116012) |
| CD45 | ALEXA-700 | Flow Cytometry | ThermoFisher (56-0451-82) |
| CD69 | PECY7 | Flow Cytometry | Biolegend (104512) |
| CD103 | FITC | Flow Cytometry | ThermoFisher (11-1031-85) |
| CCR5 | APC | Flow Cytometry | ThermoFisher (17-1951-82) |
| CXCR3 | PECY7 | Flow Cytometry | ThermoFisher (25-1831-82) |
| CD274 (PDL1) | PE | Flow Cytometry | ThermoFisher (12-5982-82) |
| CD11c | PE | Flow Cytometry | Biolegend (117308) |
| CD11b | PECy7 | Flow Cytometry | Biolegend (101216) |
| H-2Kb | PE | Flow Cytometry | Biolegend (116507) |

| | | | |
|---|---|---|---|
| H-2Kb-SIINFEKL | PE | Flow Cytometry | ThermoFisher (12-5743-82) |
| I-Ab | PERCPCY5.5 | Flow Cytometry | Biolegend (116416) |
| IFNg | PE | Flow Cytometry | Biolegend (505808) |
| Mouse IgG | HRP | Western Blot | Cell Signaling 7076S) |
| Rabbit IgG | HRP | Western Blot | Cell Signaling (7074S) |
| Rat IgG | HRP | Western Blot | Cell Signaling (7077S) |
| PD1 | None | Blocking | BioXcell (BE0273) |
| Rat IgG1a Isotype | None | Blocking | BioXcell (BE0089) |

**Supplementary Table 2. Primers**

| Primer Name | Primer Sequence (5'→3') | Purpose |
|---|---|---|
| *Mlh1* A gRNA Forward | caccgGACGGTAGTGAACCGCATAGCGG | CRISPR |
| *Mlh1* A gRNA Reverse | aaacCCGCTATGCGGTTCACTACCGTCc | CRISPR |
| *Mlh1* B gRNA Forward | caccgGGTAGTGAACCGCATAGCGGCGG | CRISPR |
| *Mlh1* B gRNA Reverse | aaacCCGCCGCTATGCGGTTCACTACCc | CRISPR |
| *Kras* A gRNA Forward | caccgGCCGCCTGCCGAATCGAGCCCGG | CRISPR |
| *Kras* A gRNA Reverse | aaacCCGGGCTCGATTCGGCAGGCGGCc | CRISPR |
| *hMLH1*_053_shRNA Forward | AATTGTGTTCTTCTTTCTCTGTATTCTCGAGAATACAGAGAAAGAAGAACACTTTTTTTAT | shRNA |
| *hMLH1*_053_shRNA Reverse | AAAAAAAGTGTTCTTCTTTCTCTGTATTCTCGAGAATACAGAGAAAGAAGAACAC | shRNA |
| *Sting*_shRNA_Forward | GCATCAAGAATCGGGTTTATT | shRNA |
| *Sting*_shRNA Reverse | CAACATTCGATTCCGAGATAT | shRNA |
| M13/pUC Forward | CCCAGTCACGACGTTGTAAAACG | Sequencing |
| M13/pUC Reverse | AGCGGATAACAATTTCACACAGG | Sequencing |
| *Mlh1* Seq Forward | GCGCGCGAATTCCCAAATCAAATGTCCGAGGGC | Sequencing |
| *Mlh1* Seq Reverse | GCGCGCGGATCCGTAGCAGGAGTTATTCGGCGT | Sequencing |
| *Ccl5* Forward | GCTGCTTTGCCTACCTCTCC | qPCR |
| *Ccl5* Reverse | TCGAGTGACAAACACGACTGC | qPCR |
| *Cxcl10* Forward | CCAAGTGCTGCCGTCATTTTC | qPCR |
| *Cxcl10* Reverse | GGCTCGCAGGGATGATTTCAA | qPCR |
| *Isg15* Forward | GGTGTCCGTGACTAACTCCAT | qPCR |
| *Isg15* Reverse | TGGAAAGGGTAAGACCGTCCT | qPCR |
| *Pro-Il18* Forward | GCCATGTCAGAAGACTCTTGCG | qPCR |
| *Pro-Il18* Reverse | TCACAGAGAGGGTCACAGCCA | qPCR |
| *Pro-Il1b* Forward | TTCAGGCAGGCAGTATCACTC | qPCR |
| *Pro-Il1b* Reverse | GAAGGTCCACGGGAAAGACAC | qPCR |
| *Gsdmd* Forward | TTCCAGTGCCTCCATGAATGT | qPCR |
| *Gsdmd* Reverse | GCTGTGGACCTCAGTGATCT | qPCR |
| *Nlrp3* Forward | ATTACCCGCCCGAGAAAGG | qPCR |
| *Nlrp3* Reverse | TCGCAGCAAAGATCCACACAG | qPCR |
| *Sting* Forward | CTACATTGGGTACTTGCGGTT | qPCR |
| *Sting* Reverse | GCACCACTGAGCATGTTGTTATG | qPCR |
| *Gapdh* Forward | CATGTTCCAGTATGACTCCA | qPCR |
| *Gapdh* Reverse | TGAAGACACCAGTAGACTCC | qPCR |
| Human *CCL5* Forward | ACCCAGCAGTCGTCTTTGTC | qPCR |

| | | |
|---|---|---|
| Human *CCL5* Reverse | CGGGTGGGGTAGGATAGTGA | qPCR |
| Human *CXCL10* Forward | AAGTGGCATTCAAGGAGTACC | qPCR |
| Human *CXCL10* Reverse | GCATCGATTTTGCTCCCCTC | qPCR |
| Human *ISG15* Forward | CGCAGATCACCCAGAAGATCG | qPCR |
| Human *ISG15* Reverse | TTCGTCGCATTTGTCCACCA | qPCR |
| Human *GAPDH* Forward | GTCTCCTCTGACTTCAACAGCG | qPCR |
| Human *GAPDH* Reverse | ACCACCCTGTTGCTGTAGCCAA | qPCR |

**Supplementary Figures**

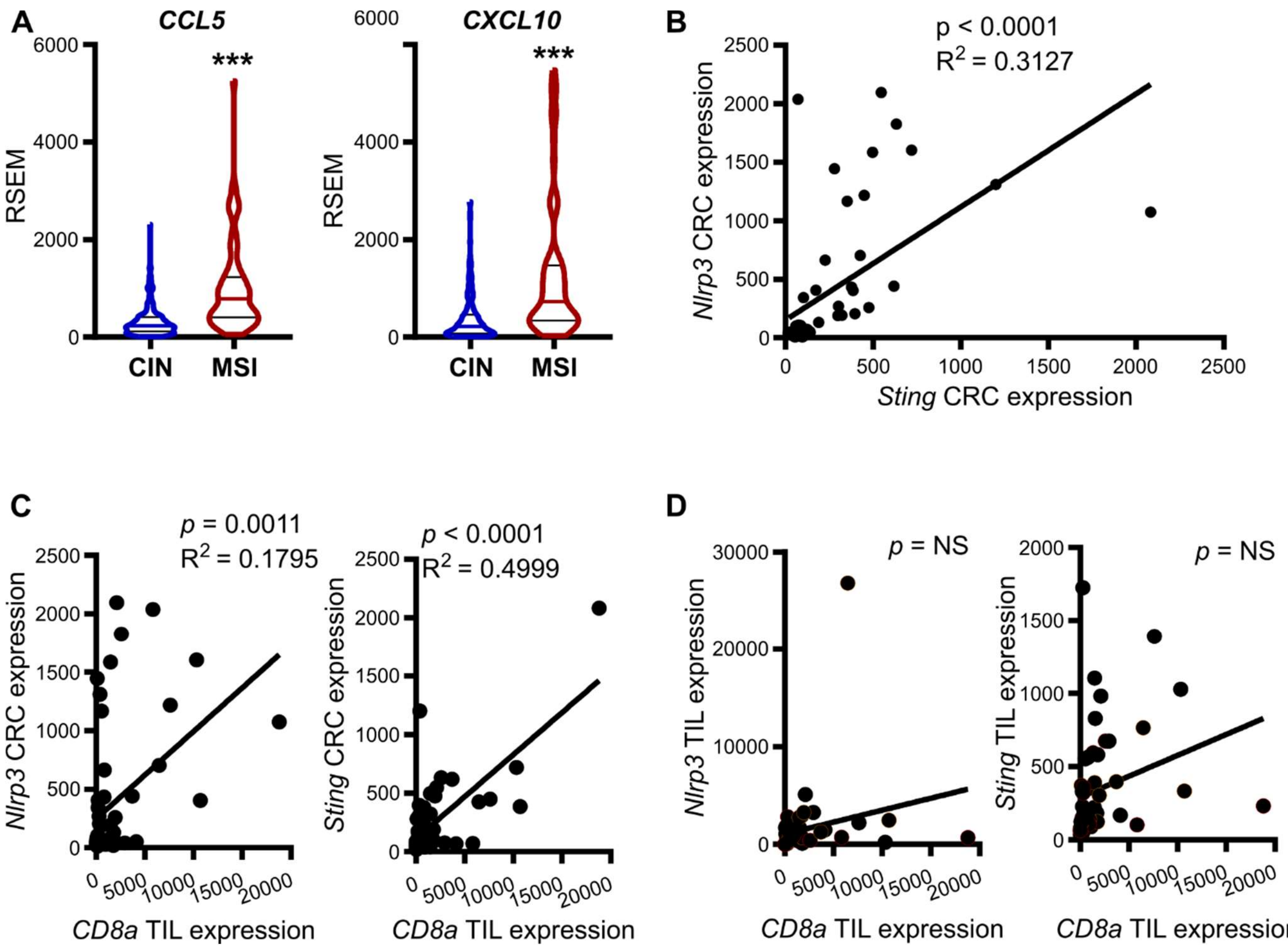


**Supplementary Figure 1. NLRP3 and STING are co-expressed in CRC cells and are correlated with CD8+ T cell infiltration. (A)** RNA expression of key cGAS/STING pathway products in human CIN and MSI colorectal cancers from the PanCancer Atlas CRC dataset on the TCGA database. Y-axis RNAseq expression is given as RNA-Seq by Expectation-Maximization (RSEM). **(B)** Correlation between the expression of indicated genes in CRC cells isolated from subcutaneously grown tumors. CRC cells were isolated and qPCR was performed on RNA extracted from these cells. **(C)** Correlation between Nlrp3 and Sting expression in CRC cells with CD8+ TIL infiltration in subcutaneously grown CRC tumors. **(D)** Lack of correlation between Nlrp3 and Sting expression in CD8+ T cells with CD8+ TIL infiltration in subcutaneously grown CRC tumors. Panels B-D show representative data from N ≥ 3 experiments. For panel A: **p < 0.001. For panels B-D: Linear regression (C.I. = 95%) with the Pearson correlation R2 and p-values reported on each graph.

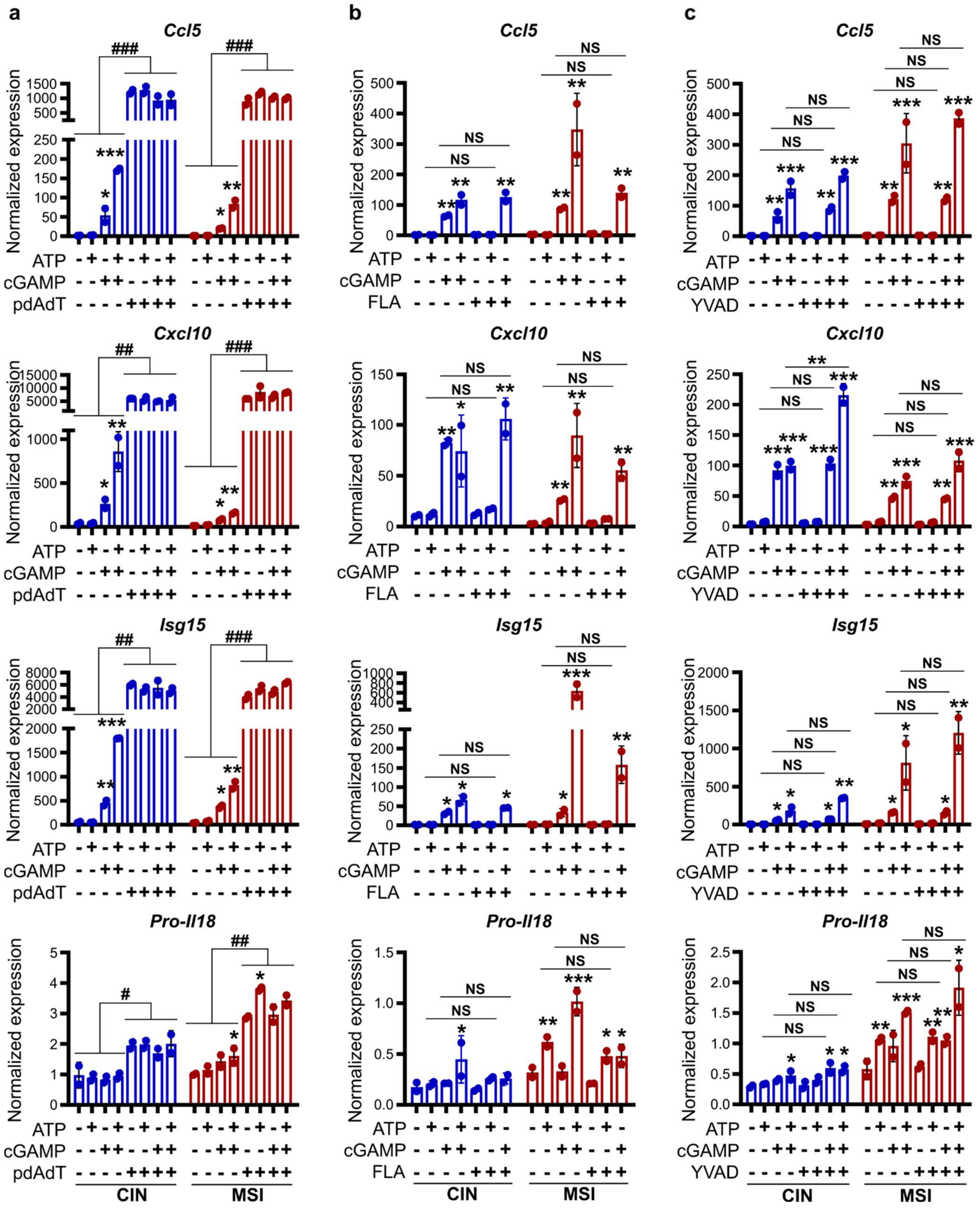


**Supplementary Figure 2. There is crosstalk between select PRRs and cGAS/STING activation but cooperative regulation between cGAS/STING and PRRs is only see for NLRP3 and is**

**independent of caspase 1.** Cells were stimulated for 24 h with 9 µg/ml 2'-3'-cGAMP and/or 2 mM ATP and addition to **(A)** 2 µg/ml poly(dA:dT) (AIM2 agonist), **(B)** 100 ng/ml flagellin (TLR5 agonist), or **(C)** 10 µg/ml YVAD (Caspase-1 inhibitor). RNA was subsequently isolated and gene expression evaluated by qPCR. All panels show representative data from N ≥ 3 experiments each with ≥ 2 biological replicates. For all panels, relative to the untreated control: *p ≤ 0.05, **p ≤ 0.01, ***p ≤ 0.001. For all panels, between indicated samples: #p ≤ 0.05, ##p ≤ 0.01, ###p ≤ 0.001.

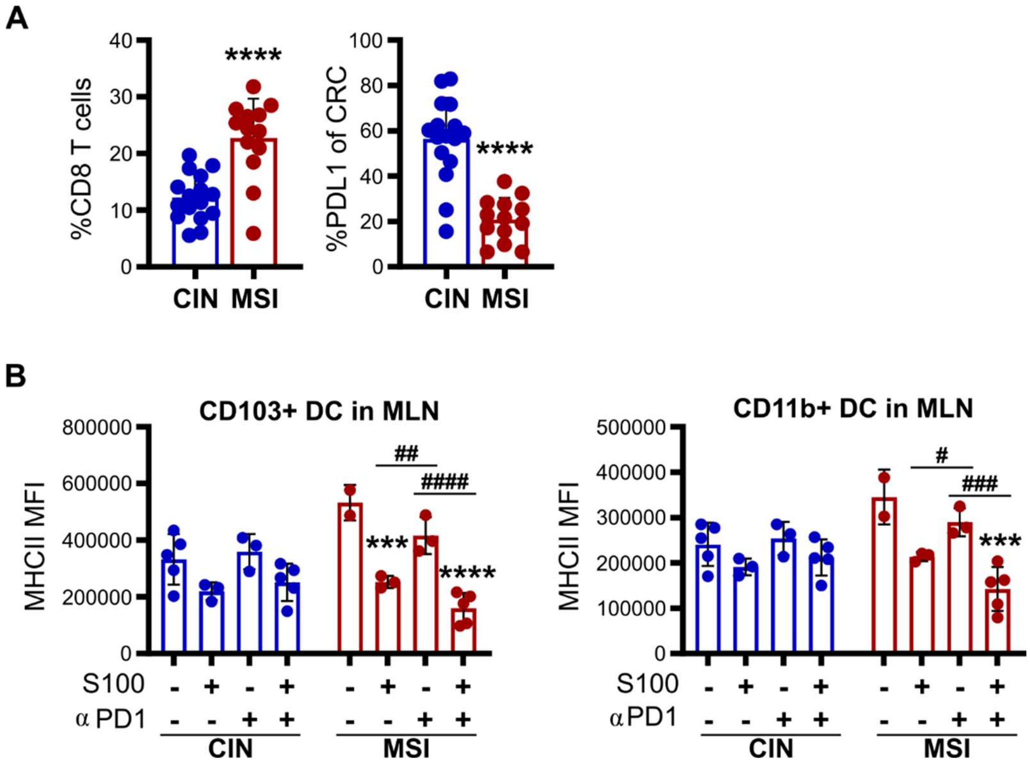


**Supplementary Figure 3. cGAS/STING and NLRP3 regulate antitumor immunity in orthotopic CRC. (A)** MC38 CRC cells were orthotopically implanted into the colons of immunocompetent mice. Mice were then treated every 3 days with 20 mg/kg MCC950 via IP injection before harvesting tumors for flow cytometric analysis. **(B)** MC38 CRC cells were orthotopically implanted into the colons of immunocompetent mice. Mice were then treated every 3 days with 1 mg/kg ADU-S100 and 200 µg of anti-PD1 or isotype via IP injection before harvesting MLNs for flow cytometric analysis. All panels show representative data from N = 3 experiments each with 3-5 mice per experiment. For all panels, relative to the untreated control: ***p ≤ 0.001, ****p ≤ 0.0001. For all panels, between indicated samples: #p ≤ 0.05, ##p ≤ 0.01, ###p ≤ 0.001.
